# Human like severe hypertriglyceridemia in a high fat fed chicken model

**DOI:** 10.1101/2024.02.19.581080

**Authors:** K. Saranya, JG Beniha, TK Deepshikha, Kriti Shankar, R Vishnu, Gopi Kadiyala, Uday Saxena

## Abstract

Cardiovascular disease (CVD) is the biggest cause of mortality globally. Controlling the risk factors for CVD such as blood cholesterol and triglycerides is the hallmark of primary prevention of CVD. There are several drugs to control cholesterol that are available but not many approaches to reducing triglycerides safely are available.

High blood triglycerides or hypertriglyceridemia in humans is classified as moderate (200-400 mg/dl plasma levels) severe (400-800 mg/dl) and very severe (>800 mg/dl). There are not many appropriate in vivo models to study human like severe hypertriglyceridemia. We show here that high fat fed chickens rapidly and in a sustained manner respond by demonstrating triglyceride levels reminiscent of severe hypertriglyceridemia in humans. Such a model could be useful in better understanding this human disease as well as serve to test new therapies.

## Introduction

Cardiovascular disease (CVD) continues to be the world’s leading cause of death. The manifestation of CVD is in the form of heart attacks, stroke, angina and myocardial ischemia. The strategy for primary prevention of CVD Involves reducing the well-established risk factors such as high blood cholesterol, type 2 diabetes, blood pressure, sedentary life style and high blood triglycerides. Hypertriglyceridemia (HTG) is a strong risk factor for cardiovascular disease. HTG has been overlooked for the last few decades because the focus of risk reduction was lowering of blood cholesterol (1,2). HTG has more recently been correlated with the incidence of stroke (3,4,5).

The usual strategy for discovery of new drugs for a disease is to first develop and test drugs in an animal model reminiscent of human conditions. These studies provide the rationale for progressing with the drug as well as show early proof of concept. In case of hypertriglyceridemia, in humans it is classified as three conditions, mild is 200-400 mg/dl, moderate is 400-800 mg/dl and severe is > 800 mg/dl of plasma triglyceride levels. In most rodent models even when fed a high fed diet, the plasma triglyceride levels go no higher than 200 mg/dl, which is considered as normal in humans. These may not be appropriate to study human like moderate or severe hypertriglyceridemia’s (6,7,8,9).

We therefore examined if the avian model may be able to achieve the levels of hypertriglyceridemia seen in human severe hypertriglyceridemia (> 800 mg/dl). The chicken models have similar complement of lipoproteins in very low-density lipoproteins (VLDL) Low density lipoproteins (LDL) and high-density lipoproteins (HDL) as humans. Similarly, the apolipoproteins such as apo B, apo C and apo A are also present. Therefore, it could a useful model to mimic severe hypertriglyceridemia seen in humans. In the present study we used the high fat fed chicken model and followed the plasma lipids, glucose and alkaline phosphatase, a biomarker for liver function. We observe that high fat diet rapidly induces hypertriglyceridemia which is consistently above 800 mg/dl reminiscent of human severe hypertriglyceridemia. Thus, such a model could be used to rapidly test drugs for treatment of human severe hypertriglyceridemia.

## Methods

- Selected 12 healthy chickens n= 6 (Giriraja species). A leg tag is attached for numbering them
- Feed Preparation:
- Breeder Mash Feed+ 20% Pure coconut oil
- 150g feed /bird – per day
- Blood collection:
- The site of blood collection is wiped with 70% ethanol
- The required volume of blood from is drawn brachial wing vein
- The plasma is analyzed for the following blood parameters
- Glucose
- Cholesterol
- Triglycerides
- Alkaline Phosphatase as marker for liver dysfunction

## Results

### Study design

Shown below in Figure 1 is the timeline of high fat diet initiation and blood collection for measuring various blood parameters.

**Figure 1.**
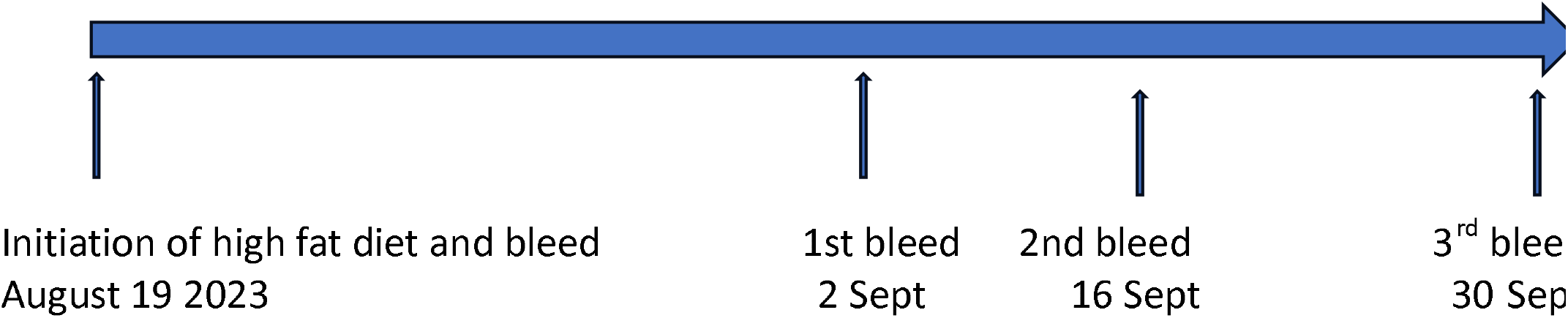

#### Various plasma parameters after diet initiation

Shown below are the various plasma parameters after initiation of high fat diet in the chickens.

### Triglycerides

As shown below figure 2, there was rapid induction of hypertriglyceridemia with the high fat diet. The individual animal plasma triglyceride levels are shown below. In most animals, week on week, for the duration of study the levels averaged over 800 mg/dl suggesting a human like severe hypertriglyceridemia profile. As mentioned before such levels reflect human disease like phenotype.

**Figure 2:**
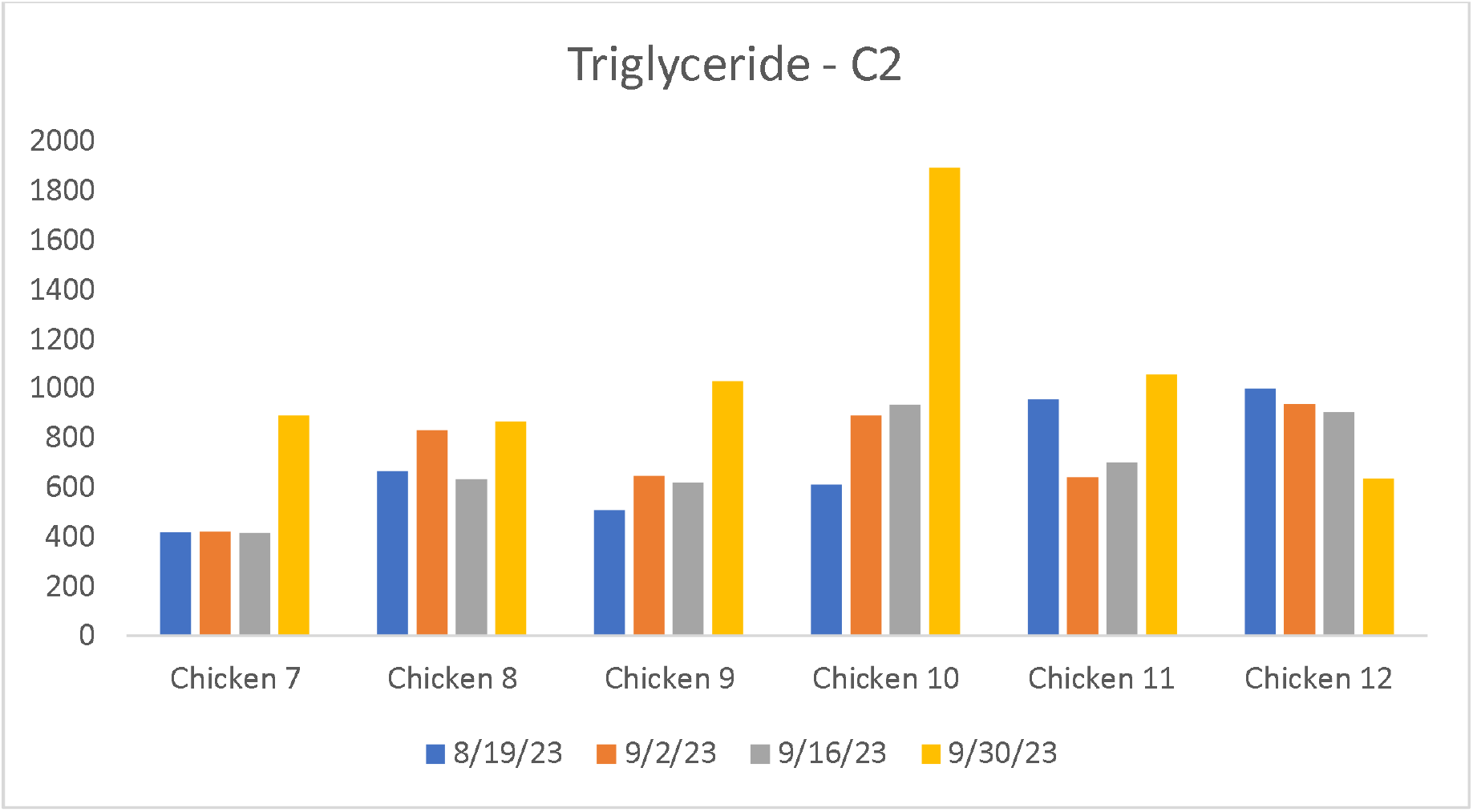
Triglyceride values from individual birds at different time points

### Plasma Glucose

We also looked at the plasma glucose levels. As shown in figure 3 the glucose levels of around >200 mg/dl approximated a moderate diabetic condition in humans in all animals thru duration of study. This is an additional benefit of the model in that this model may be potentially useful to study moderate diabetes.

**Figure 3:**
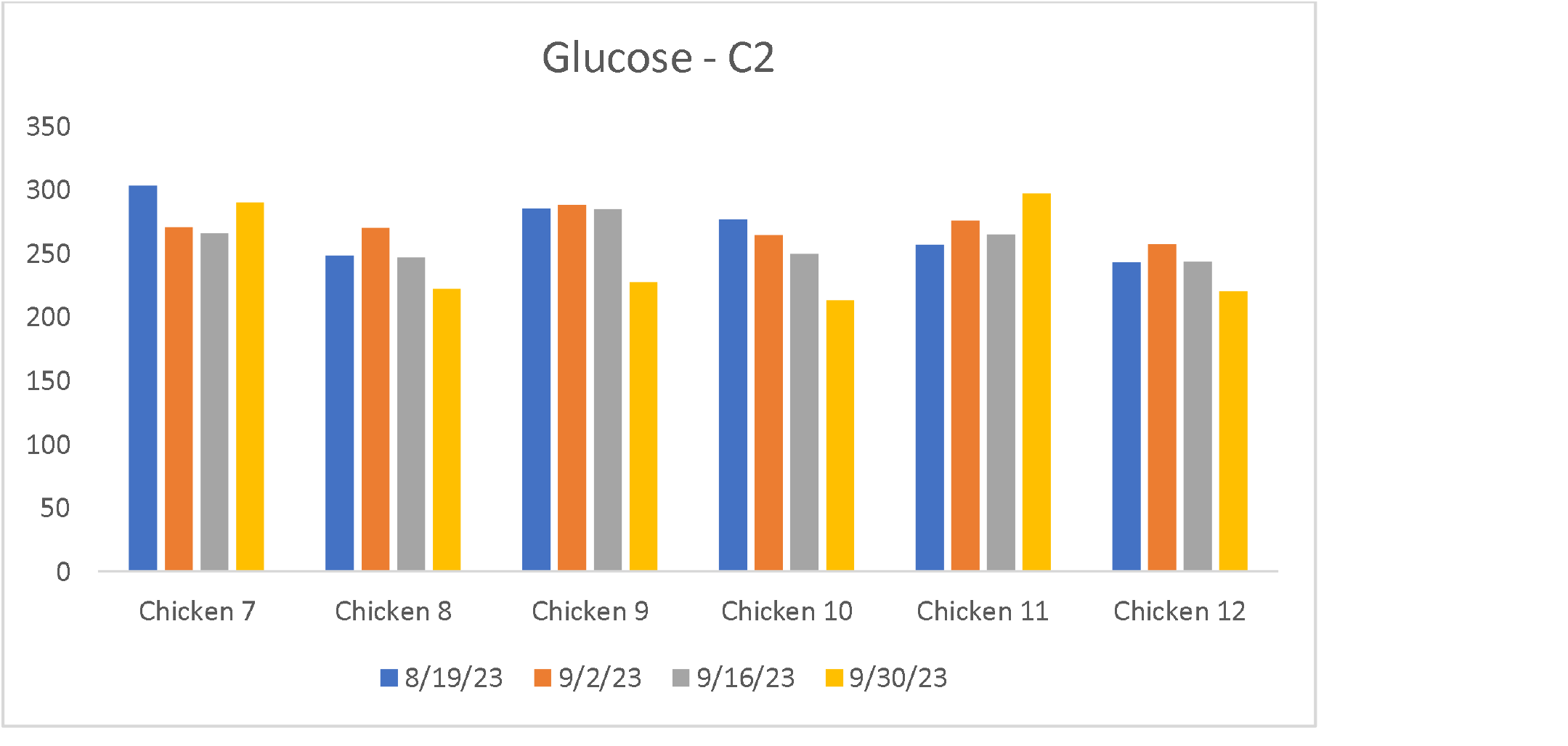
Glucose values from individual birds at different time points

### Plasma cholesterol

The levels of plasma cholesterol were also examined. As shown in figure 4, the cholesterol levels in individual animals varied with lowest of 75 mg/dl to high of >250 mg/dl. There was individual variation and because of which it may difficult to use this model to study hypercholesterolemia in human where the levels range 200-400 mg/dl.

**Figure 4:**
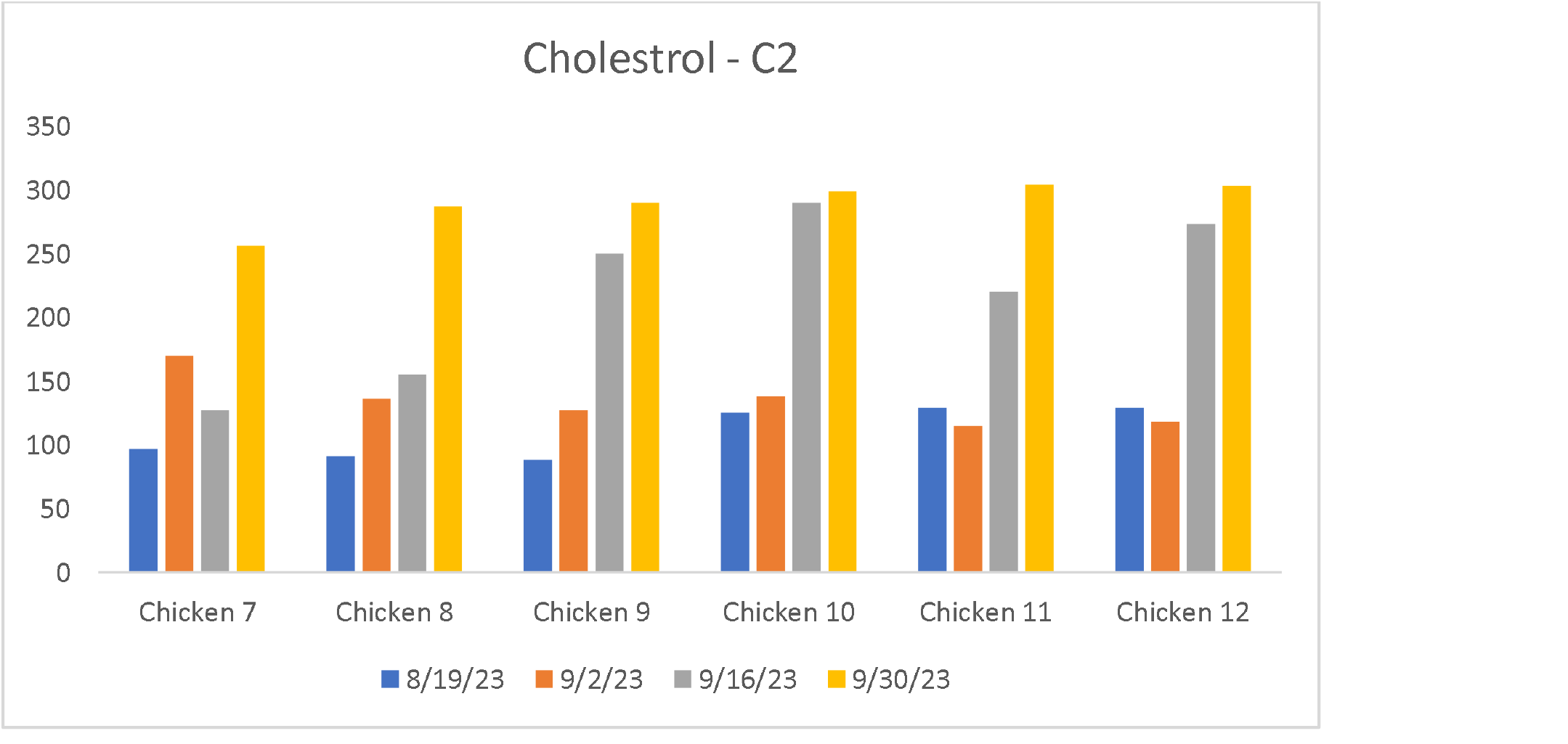
Cholesterol values from individual birds at different time points

### Alkaline phosphatase levels

Plasma alkaline phosphatase levels are often used as a biomarker for fatty liver or other liver dysfunctions. We monitored the levels of this enzyme in our study to see if the levels were high. As shown in figure 5, ALP levels were high relative to a human level, likely due to the animals being fed a high fat diet which may induce fatty liver.

**Figure 5:**
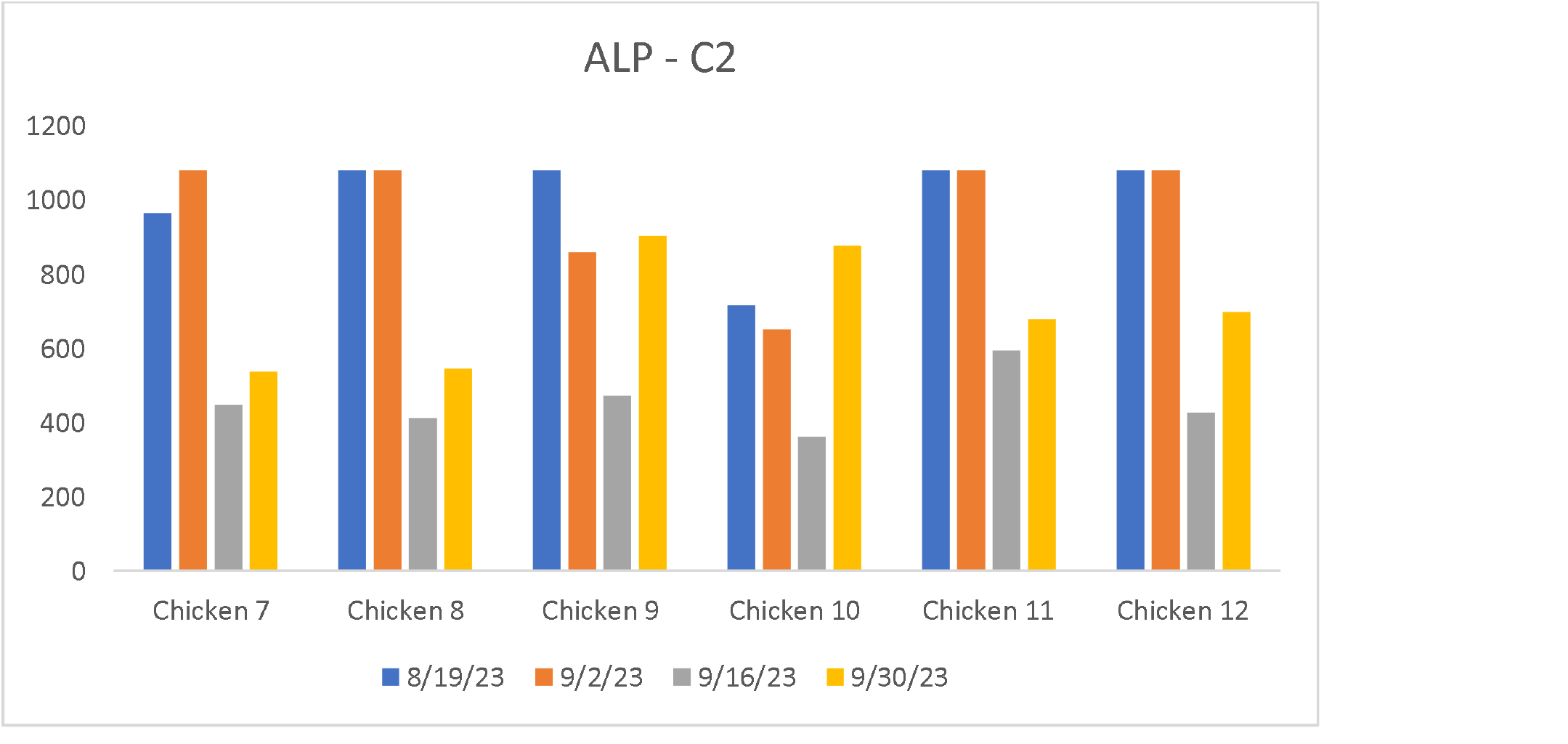
ALP values from individual birds at different time points

### Lipoprotein distribution of plasma cholesterol

The distribution of total plasma cholesterol in various lipoprotein fractions was also determined. As shown in figure 6 below, the bulk of cholesterol was found in the VLDL (very low-density lipoprotein) fraction in all animals. This partitioning of plasma cholesterol mostly in VLDL is typical seen during hypertriglyceridemia, VLDL is the major and most abundant lipoprotein during hypertriglyceridemia is similar to the human condition.

**Figure 6:**
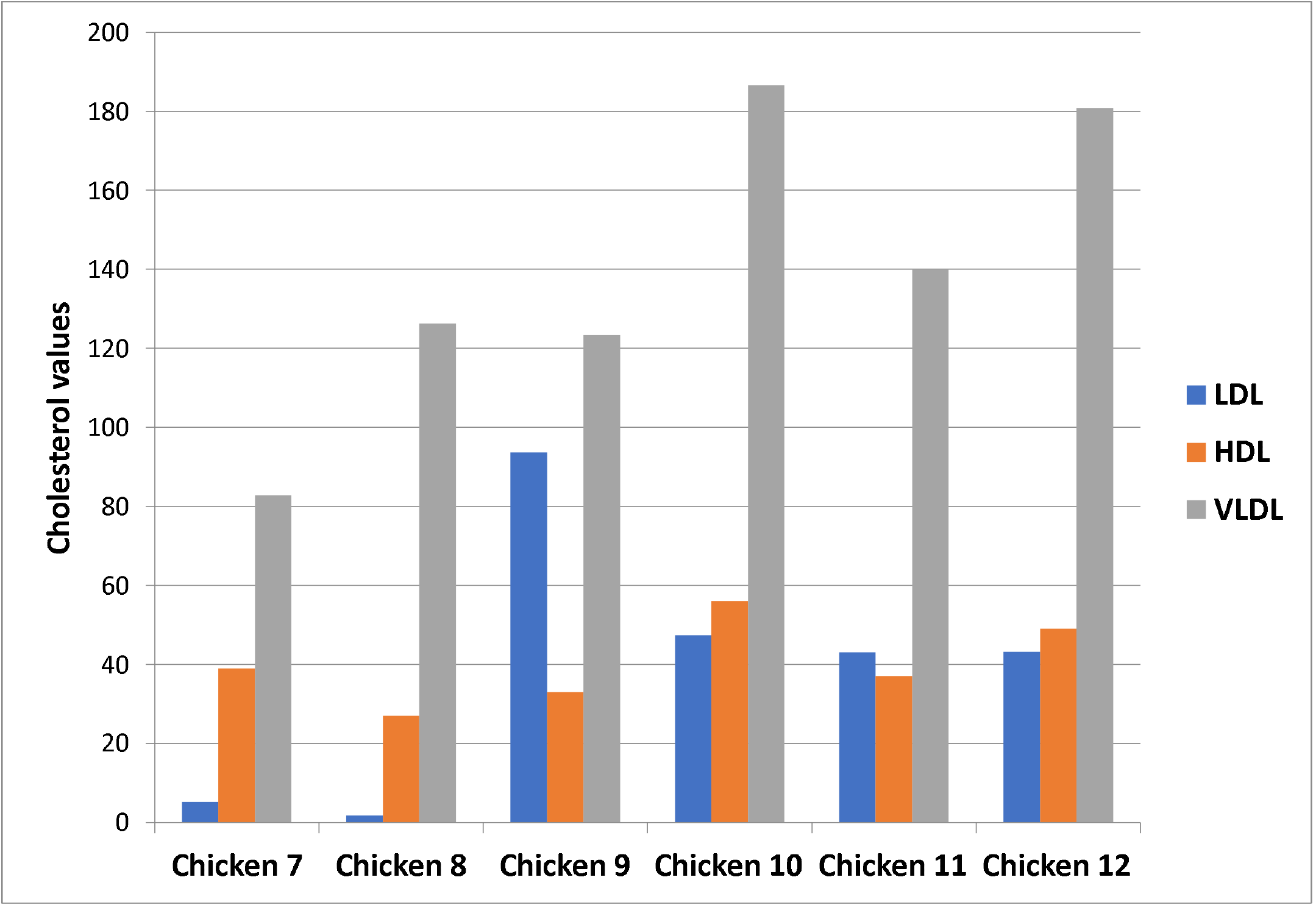
Lipoprotein cholesterol distribution in various lipoproteins

### Comparison with human values

The average values of parameters of all animals are shown below in figure 7 with a comparison with human phenotype. Triglycerides were clearly like severe human levels, while cholesterol and glucose approached human disease levels as did ALP.

**Figure 7:**
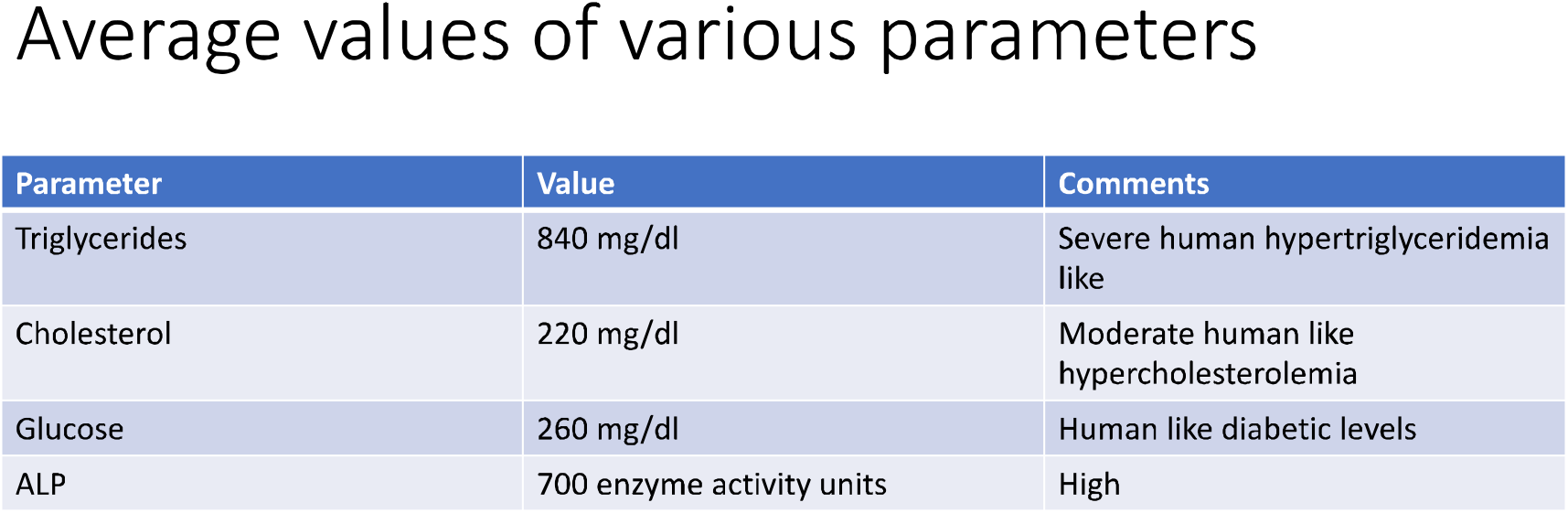
Comparison of chicken values with human

## Discussion

Dyslipidemia in humans continues to be a major problem and can result in CVD. Most attention till date was focused on high plasma cholesterol as the risk factor to target. But now triglycerides have become am emerging risk factor for CVD. With regards to animal models to study human like hypertriglyceridemia the existing rodent models do not present with triglyceride levels which mimic human severe hypertriglyceridemia.

To this end we explored a high fat chicken as a model to produce human like hypertriglyceridemia. We show here that in this model high fat diet is able to induce triglyceride levels reminiscent of human like severe hypertriglyceridemia. In addition, other plasma parameters associated with metabolic syndrome, cholesterol and glucose also approached human disease like levels.

The chicken as a model for dyslipidemia is largely unexplored with very few studies published. We were especially very interested to find that the model shows several features seen in human metabolic syndrome. We propose that fat fed chicken can be an in-expensive and easy to use with wide availability model to understand and test drugs for human severe hypertriglyceridemia.

